# Eifficient Mitigation of Copper Induced Cellular Dysfunction Using Chitosan Based Iron Oxide Nanoparticles

**DOI:** 10.64898/2026.08.24.746706

**Authors:** Sakshi Chouhan, Shilpa Chandra, Chayan Kanti Nandi

## Abstract

Copper (Cu^2+^) is an essential redox-active micronutrient, but agricultural soils are increasingly contaminated by Cu^2+^ from mining, industrial discharge, and intensive agrochemical use, pushing concentrations beyond levels plants can tolerate. Excess Cu^2+^ triggers Fenton-like reactive oxygen species (ROS) generation, mitochondrial dysfunction, and impaired growth. Existing mitigation strategies, such as soil amendments, phytoremediation, antioxidants, and different chelators have been explored to reduce Cu^2+^ toxicity, but their effectiveness can be limited by immobilization, poor specificity, and environmental persistence. The present work introduces a nanoparticle-based strategy for the direct sequestration of excess Cu^2+^, coupled with protection against the oxidative damage caused by Cu^2+^ stress. Here, we report MPA-iron oxide nanoparticles (MIONPs), sequentially functionalized with chitosan, glutathione, and 3-mercaptopropionic acid, designed to simultaneously scavenge ROS, restore redox homeostasis, and chelate Cu^2+^via surface thiol groups. MIONPs showed a significant increase in Cu^2+ 2+^-binding capacity over bare iron oxide nanoparticles (BIONPs) and, in Cu^2+^ -stressed *Solanum lycopersicum* seedlings, significantly improved germination and root/shoot growth, reduced intracellular ROS, restored mitochondrial membrane potential, and preserved nuclear integrity. This integrated design establishes MIONPs as a promising, dual-function nanoplatform for sustainable Cu^2+^ stress management in agriculture.

## Introduction

Copper (Cu^2+^) is an essential redox-active micronutrient that supports photosynthetic electron transport, mitochondrial respiration, antioxidant defense, and cell wall lignification in plants^1,2^. However, its biological indispensability is bounded by a narrow concentration window, once soil Cu^2+^ levels exceed this threshold, the same redox chemistry that makes Cu^2+^ essentially becomes the source of cellular damage^3,4^. Over the past few decades, mining runoff, industrial discharge, and the intensive use of Cu^2+^-based agrochemicals such as fungicides have driven a steady and, in many regions, alarming rise in soil Cu^2+^ concentrations^5,6^. This escalating contamination has become a major constraint on crop productivity and soil health worldwide, motivating an urgent need for effective, targeted mitigation strategies.

At the physiological level, excess Cu^2+^ promotes Fenton-like reactions that generate reactive oxygen species (ROS), overwhelming the plant’s endogenous antioxidant machinery^4,7,8^. This oxidative stress disrupts mitochondrial membrane integrity and impairs photosynthetic efficiency, and the resulting mitochondrial dysfunction has increasingly been linked to retrograde signaling that propagates stress signals toward the nucleus, ultimately compromising chromatin organization and cell viability^9,10^. A range of strategies has been developed to counter Cu^2+^ phytotoxicity, spanning soil-level interventions such as biochar, lime, and zeolite amendments to immobilize Cu^2+^ phytoremediation using Cu^2+^ - tolerant plant species, exogenous antioxidant supplementation with glutathione (GSH) or ascorbic acid^11^; synthetic chelating agents such as EDTA^12^; and, more recently, engineered nanomaterials^13–15^. Among nanomaterial-based approaches, ZnO nanoparticles suppress Cu^2+^ -induced ROS accumulation^16,17^, while SiO_2_ nanoparticles restrict Cu^2+^ uptake by strengthening the plant cell wall^18^. Selenium nanoparticles enhance antioxidant metabolism and maintain cellular redox homeostasis^13,19^. ZnO and TiO_2_ nanoparticles may also exhibit dose-dependent phytotoxicity^20–22^, whereas selenium nanoparticles have a relatively narrow beneficial-to-toxic concentration range^23,24^. However, these materials mainly works on the oxidative consequences of Cu^2+^ stress or reduce Cu^2+^ uptake, with limited ability to address the accumulated Cu^2+^ within plant tissues.

Iron oxide (Fe3O4) nanoparticles offer a distinct advantage in this context owing to their intrinsic nanozyme activity peroxidase- and catalase-mimetic behavior that enables direct enzymatic breakdown of ROS^25,26^. Yet bare Fe3O4 nanoparticles (BIONPs) are limited by poor colloidal stability^27^ and, more importantly, possess no inherent Cu^2+ 2+^-binding functionality, leaving the metal-sequestration side of the problem entirely unaddressed. Various surface-engineering strategies have since been explored to improve the physicochemical performance of Fe3O4 nanoparticles^28^, but these efforts have largely optimized isolated properties such as stability or biocompatibility rather than building an integrated system capable of tackling both ROS accumulation and Cu^2+^toxicity together. As a result, Fe3O4-based nanoparticles purpose-built to simultaneously regulate Cu^2+^accumulation and Cu^2+^ -induced oxidative stress remain notably scarce.

To address this gap, we designed MPA-iron oxide nanoparticles (MIONPs) through sequential surface functionalization of BIONPs with chitosan, glutathione (GSH), and 3-mercaptopropionic acid (MPA), using the chitosan/GSH-coated particle as an intermediate^29,30^. Unlike prior single-mechanism nanomaterials, this rational, layer-by-layer design integrates three complementary functions within one platform. The intrinsic nanozyme activity of the Fe3O4 core, GSH-mediated intracellular antioxidant reinforcement, and selective Cu^2+^chelation via surface thiol groups introduced by MPA alongside markedly improved colloidal stability. In this study, we evaluated the potential of MIONPs for managing excess Cu^2+^in plants and Cu^2+^ -contaminated medium. The efficacy of MIONPs was evaluated in *Solanum lycopersicum*at physiological and cellular levels. At the physiological level, Cu^2+^ stress and MIONP-mediated recovery were assessed by monitoring germination, root development, and shoot growth. At the cellular level, the response was examined along the Cu^2+^ -induced stress cascade, beginning with intracellular ROS accumulation as an early indicator of oxidative stress, followed by mitochondrial dysfunction and nuclear alterations. Mitochondria were examined because of their central role in cellular energy production and their susceptibility to Cu^2+^ -induced oxidative damage, while the nucleus was evaluated as a key site of downstream cellular damage and chromatin organization. In parallel, the Cu^2+^ binding capacity of MIONPs was assessed to determine their ability to capture excess Cu^2+^ ions. This work establishes MIONPs as a mechanistically grounded, dual-function nanoplatform with clear potential for sustainable agricultural applications, offering a targeted route to managing Cu^2+^-contaminated soils without the trade-offs that limit existing single-mechanism mitigation strategies.

## Results and discussions

### Characterization of BIONPs, Intermediate Nanocomposite and MIONPs

As outlined in Scheme 1, the stepwise surface functionalization of BIONPs into the chitosan/GSH-coated intermediate and finally into MIONPs was undertaken to progressively introduce GSH-mediated antioxidant capacity and MPA-derived thiol groups onto the Fe3O4 nanozyme core. The successful synthesis and sequential surface functionalization of BIONPs into the intermediate nanocomposite and finally MIONPs were confirmed using complementary physicochemical characterization techniques. TEM images **(Figure 1a-c)** revealed that BIONPs possessed a nearly spherical morphology with an average particle size of 8.6 ± 2.8 nm. After coating with chitosan and GSH, the particle size is 9.7 ± 1.2 nm; subsequent conjugation with MPA further increased the particle size to 10.52 ± 0.94 nm, of MIONPs **(Figure 1d-f)**^**31**^. HR-TEM images **(Figure 1g-i)** further revealed lattice fringes with d-spacings of 0.259-0.262 nm, corresponding to the (311) crystal plane of magnetite, indicating that the crystalline Fe3O4 core remained intact throughout functionalization^32^. SEM-EDX analysis **(Supplementary Figure S1)** further verified the sequential surface modification: BIONPs exhibited only Fe (23.63%) and O (76.37%), whereas the intermediate nanocomposite showed additional C (8.01%), N (6.13%), and S (0.47%) signals with a reduction in Fe content to 19.73%. After MPA conjugation, the carbon and sulfur contents further increased to 14.89% and 0.94%, respectively, while Fe decreased to 17.68%, confirming surface coverage by the organic coating.

**Figure 1.**
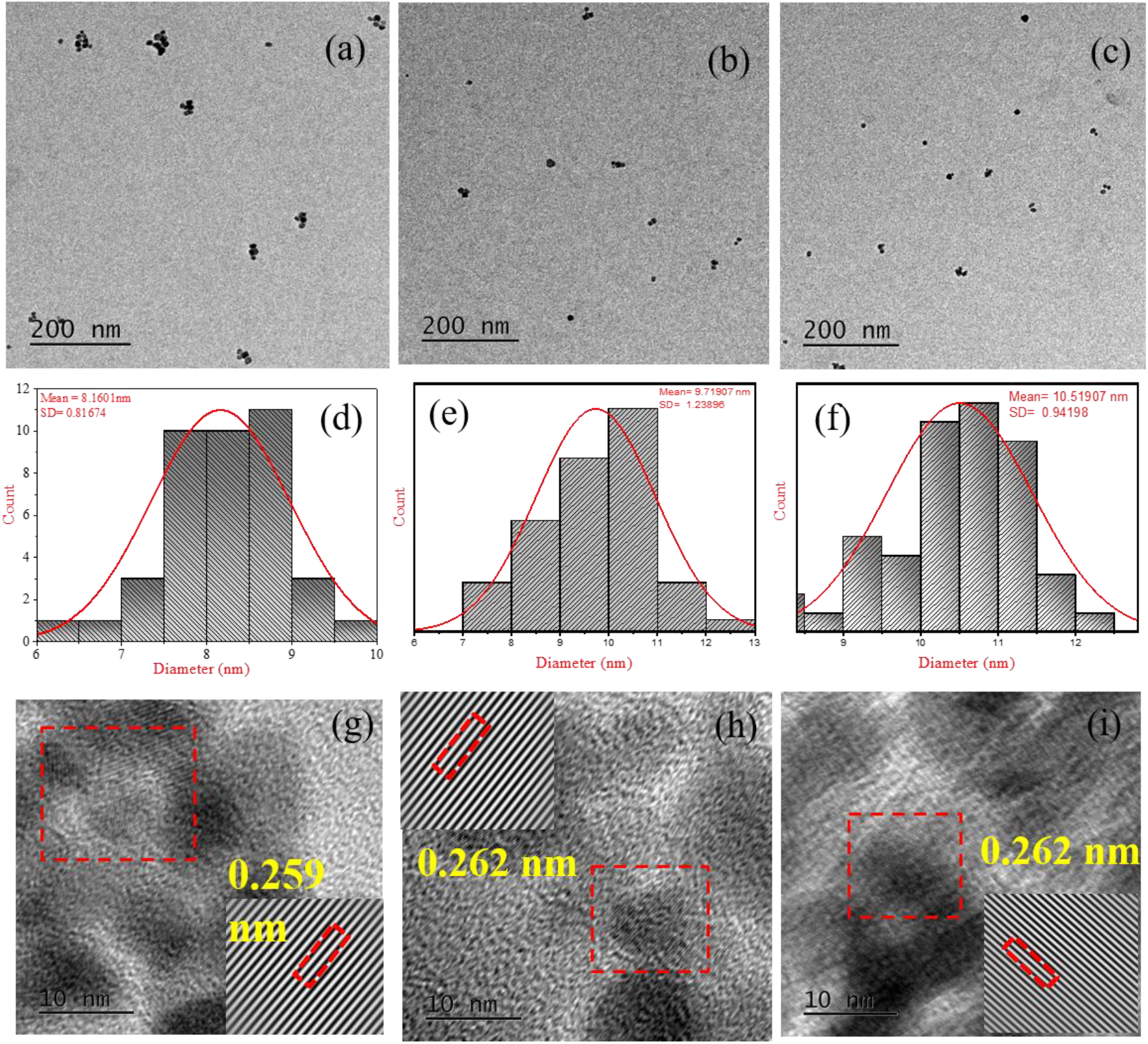
TEM characterization of stepwise functionalized nanoparticles. (a-c) TEM images of BIONPs, the intermediate, and MIONPs, respectively. (d-f) Particle size distribution histograms derived from the TEM images of BIONPs, the intermediate, and MIONPs, respectively. (g-i) HR-TEM images showing lattice fringes with corresponding d-spacing values of (g) 0.259 nm for BIONPs, (h) 0.262 nm for the intermediate, and (i) 0.262 nm for MIONPs.

The chemical functionalization of the nanoparticles was investigated by FTIR spectroscopy **(Figure 2a)**. BIONPs displayed the characteristic Fe-O stretching vibration at 554 cm^−1^, while chitosan exhibited characteristic bands at 1047 cm^−1^ (C–O–C glycosidic stretch), 1655 cm^−1^ (amide I/N–H bending), and 2865/2988 cm^−1^ (C–H stretching). In the final MIONPs, complete disappearance of the MPA carboxylic acid band at 1695 cm^−1^ together with the appearance of a new band at 1168 cm^−1^ and 1220 cm^−1^, assigned to C-N stretching, confirmed successful amide bond formation between the carboxyl group of MPA and the amino groups of chitosan through EDC/NHS coupling chemistry^33–36^.

**Figure 2.**
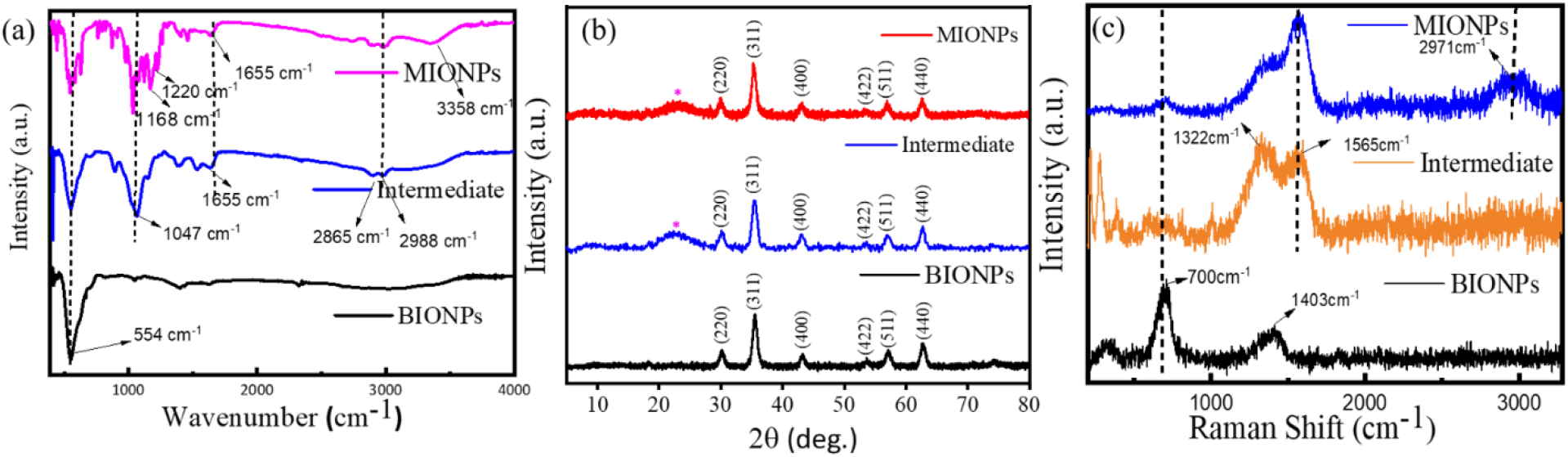
Surface and chemical characterization of stepwise functionalized nanoparticles. (a) FTIR spectra of BIONPs, the intermediate, and MIONPs, confirming the presence of characteristic functional groups. (b) XRD patterns of BIONPs, the intermediate, and MIONPs, confirming retention of the crystalline magnetite phase across functionalization. (c) Raman spectra of BIONPs, the intermediate, and MIONPs, confirming the phase stability of the magnetite core

The crystalline structure of the nanoparticles was examined by XRD **(Figure 2b)**, which showed the characteristic diffraction peaks of inverse spinel magnetite at 35.54°, 57.14°, and 62.72°, corresponding to the (311), (511), and (440) planes. Retention of these peaks, together with the appearance of a broad peak at 22.81° attributed to chitosan, confirms successful surface functionalization while preserving the crystalline Fe3O4 core. Raman spectra (Figure 2c) further supported these observations by retaining the characteristic A1g vibration of magnetite at 700 cm^−1^ in all samples. The appearance of new bands at 1322 cm^−1^ (amide III) and 1565 cm^−1^ (amide II) after chitosan/GSH coating confirmed successful GSH incorporation, whereas the emergence of an additional peak at 2971 cm^−1^ in MIONPs shows MPA conjugation^37^.

The surface chemical composition was further analyzed by XPS **(Figure 3a-c)**. The survey spectrum of BIONPs contained only Fe (28.99%) and O (71.01%), confirming the formation of pure magnetite nanoparticles. After chitosan/GSH coating, the intermediate nanocomposite exhibited C1s (45.17%), N1s (6.60%), O1s (43.83%), Fe2p (3.84%), and S2p (0.56%) signals, confirming successful surface functionalization. Following MPA conjugation, the carbon and sulfur contents further increased to 57.33% and 0.71%, respectively, while Fe decreased to 1.70%, indicating greater organic surface coverage. High-resolution Fe2p spectra **(Supplementary Figure S2a-c)** confirmed the coexistence of Fe^2+^ and Fe_3+_, demonstrating that the magnetite core remained chemically stable during functionalization. Deconvoluted S2p spectra (Figure 3f-g) showed characteristic peaks at 161.0 and 162.1 eV for the intermediate and 161.7 and 162.9 eV for MIONPs, confirming the presence of free thiol (-SH) groups. Deconvoluted C1s spectra **(Figure 3d-e)** showed an increased contribution at 286 eV, together with new N1s peaks at 398 and 400 eV **(Supplementary Figure S2g-h)**, confirming successful amide bond formation after MPA conjugation. The preservation of free thiol groups after conjugation provides abundant coordination sites for efficient Cu^2+^ binding^38–41^.

**Figure 3.**
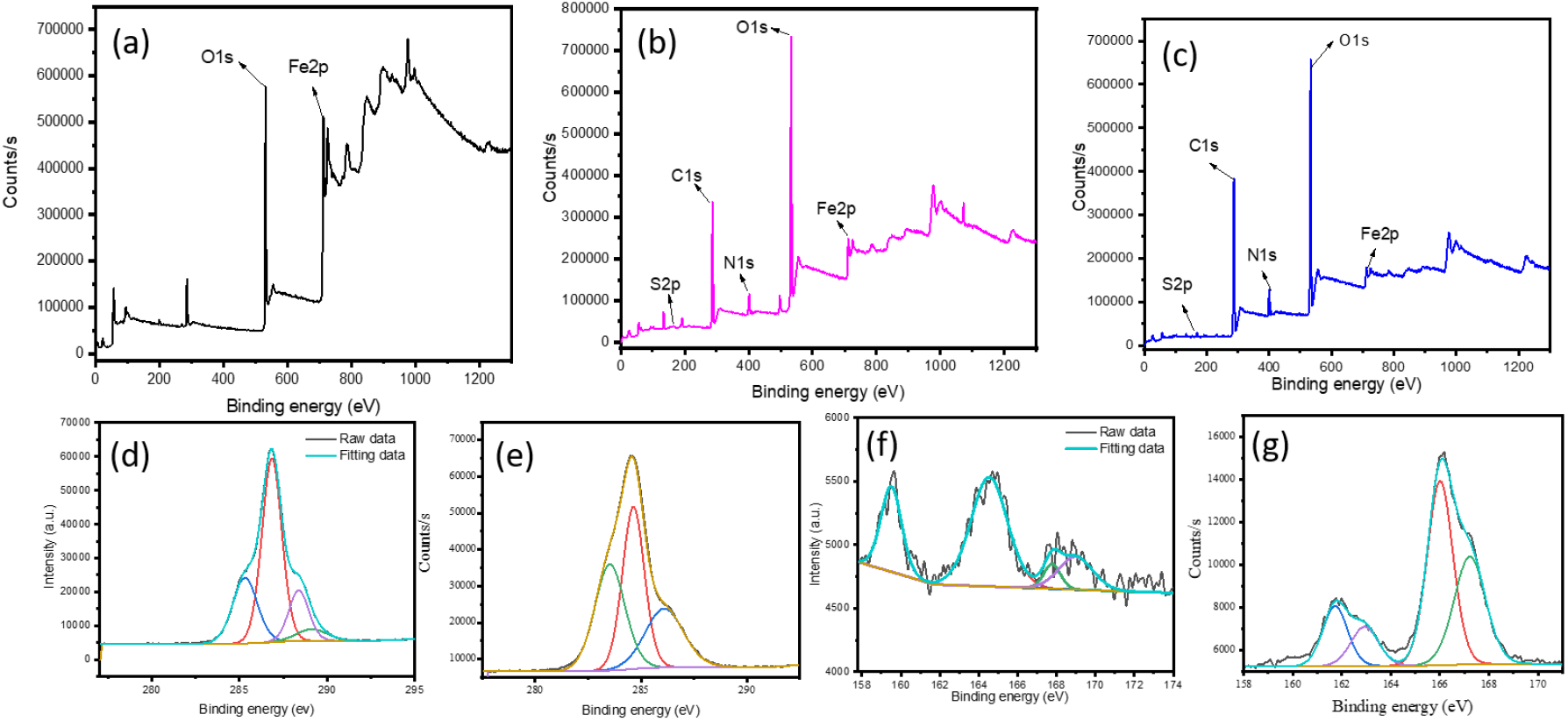
XPS data. (a) XPS survey spectrum of BIONPs. (b) XPS survey spectrum of the intermediate (Fe3O4/Chitosan/GSH). (c) XPS survey spectrum of MIONPs . (d-e) Deconvoluted C1s spectra of the intermediate and MIONPs, respectively. (f-g) Deconvoluted S2p spectra of the intermediate and MIONPs, respectively.

The colloidal properties of the nanoparticles were evaluated by zeta potential and DLS measurements. Zeta potential (Supplementary Figure S3a) increased progressively from +3.61 mV for BIONPs to +8.36 mV for the intermediate nanocomposite and finally to +35.3 mV for MIONPs, indicating a substantial improvement in colloidal stability following complete surface functionalization^42^. DLS analysis **(Supplementary Figure S4)** showed hydrodynamic diameters of 97.01 nm (PDI 0.687) for BIONPs, 85 nm (PDI 0.752) for the intermediate nanocomposite, and 95.23 nm (PDI 0.590) for MIONPs; the larger hydrodynamic sizes relative to TEM are attributed to the hydrated polymer shell surrounding the nanoparticles in aqueous suspension. The progressive deposition of the organic shell was further supported by AAS analysis **(Supplementary Figure S3b)**, which showed a gradual reduction in iron content to 67.10% in the intermediate nanocomposite and 52.04% in MIONPs^43^. ICP-MS analysis (Supplementary Figure S3c) demonstrated enhancement in Cu^2+^ adsorption efficiency, increasing from 11.78% (Q_e_ = 0.589 µg g^−1^) for BIONPs to 22.29% (Q_e_ = 1.115 µg g^−1^) for the intermediate nanocomposite and 47.61% (Q_e_ = 2.381 µg g^−1^) for MIONPs^43–45^. This about four-fold improvement is attributed to the synergistic effect of GSH and the additional surface thiol groups introduced by MPA, which strongly coordinate Cu^2+^ through soft acid-soft base interactions. Collectively, these characterization results confirm the successful fabrication of MIONPs with preserved magnetic crystallinity, efficient surface functionalization, enhanced colloidal stability, and efficient Cu^2+^ binding capability.

### Biological Evaluation of MIONPs under Copper Stress

To evaluate the protective efficacy of the developed nanoplatform against Cu^2+^ -induced phytotoxicity, a systematic physiological and cellular assessment was performed in *Solanum lycopersicum* seedlings. Seedlings were subjected to stress and subsequently treated with different nanoparticle formulations to compare their ability to alleviate Cu^2+^ -induced damage. To determine whether MIONPs can alleviate Cu^2+^ induced phytotoxicity, seedlings were exposed to 100 and 200 µM Cu^2+^, selected to represent moderate and relatively severe Cu^2+^ stress conditions, respectively. Following Cu^2+^ exposure, seedlings were treated with BIONPs, intermediate nanocomposite and MIONPs, while untreated seedlings served as the control. BIONPs and the intermediate formulation were included as comparative materials to evaluate the contribution of each successive surface modification and determine the enhanced efficacy of MIONPs. Physiological parameters were evaluated to determine Cu^2+^ induced growth inhibition and treatment-mediated recovery, followed by analysis of intracellular ROS accumulation, mitochondrial membrane potential, and nuclear chromatin organization to elucidate the progression of Cu^2+^ -induced cellular damage and the protective response mediated by MIONPs.

To assess the physiological effects of Cu^2+^ stress and the protective efficacy of MIONPs, *Solanum lycopersicum* seedlings were exposed to 100 and 200 µM Cu^***2+***^ **(Figures 4b, c)** and treated with BIONPs (**Figures 4d, g)**, the intermediate (**Figures 4e, h)** and MIONPs (**Figures 4f, i)**. Germination percentage (GP), germination rate (GR), synchronization index (SI), and mean germination time (MGT) were evaluated to assess the effects of Cu^2+^ stress on the extent, rate, uniformity, and timing of seed germination, respectively^46^, while root and shoot development were assessed to determine the impact of Cu^2+^ stress on seedling growth **(Figure 5)**. Excess Cu^2+^ disrupts plant physiology by interfering with enzyme activity, nutrient uptake, and cellular metabolism. It promotes excessive ROS accumulation, causing oxidative stress and damage to cellular components. Cu^2+^ toxicity also impairs membrane and mitochondrial functions, reducing cellular energy production. These disturbances ultimately restrict cell division, cell elongation, and root and shoot development. As these physiological parameters reflect the integrated phenotypic response of seedlings to Cu stress.

**Figure 4.**
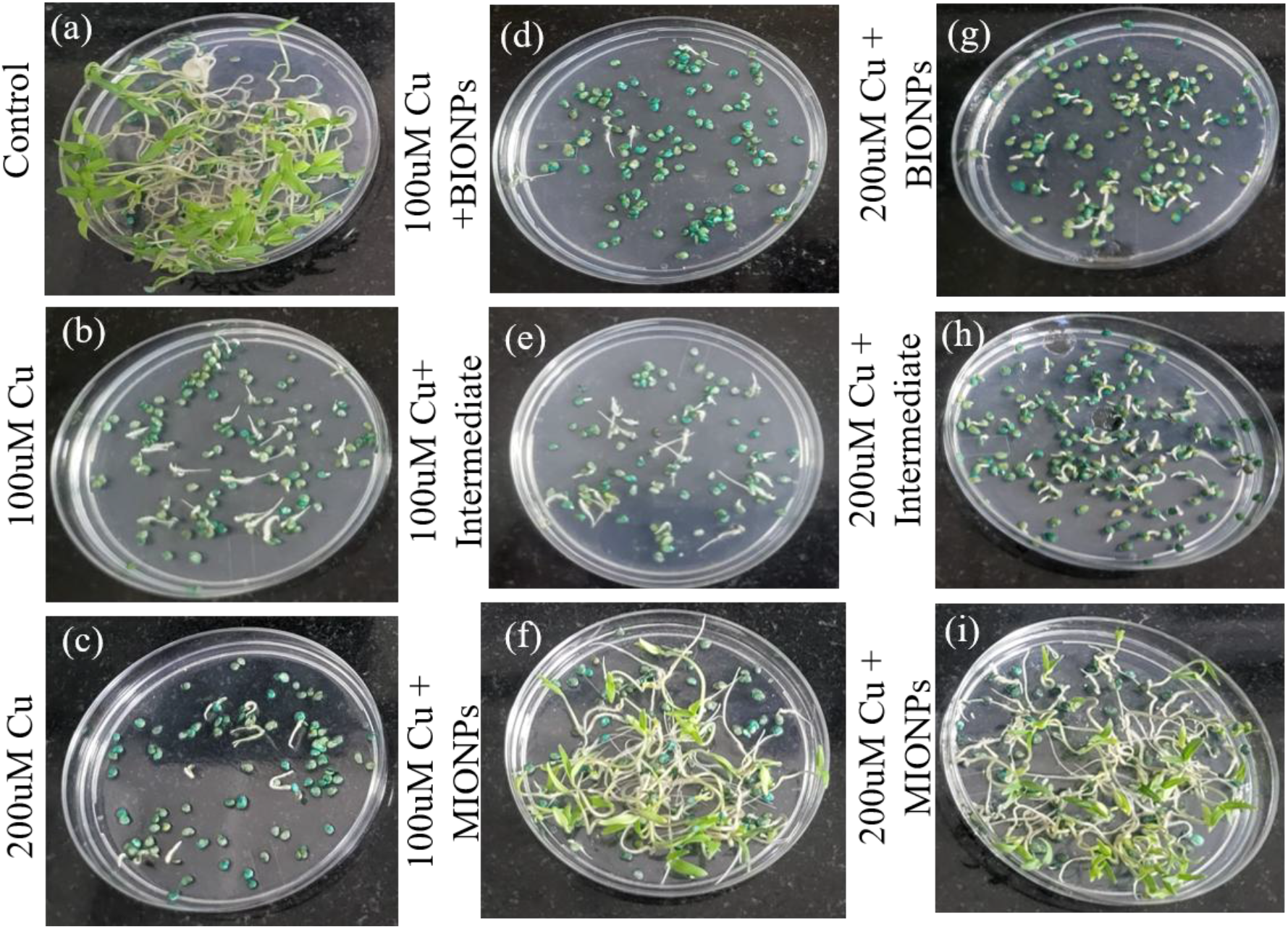
Physiological response of Solanum lycopersicum seedlings under copper stress and nanoparticle treatment (15 days). (a) Control showing normal germination and healthy root-shoot development. (b, c) Seedlings exposed to 100 µM and 200 µM Cu^2+^, respectively, showing reduced germination and inhibited growth. (d, g) Cu^2+^-stressed seedlings (100 and 200 µM, respectively) treated with BIONPs showing slight improvement in growth. (e, h) Cu^2+^-stressed seedlings treated with Intermediate showing moderate recovery in germination and growth. (f, i) Cu^2+^-stressed seedlings treated with MIONPs showing significant restoration of root and shoot growth, comparable to control conditions.

**Figure 5.**
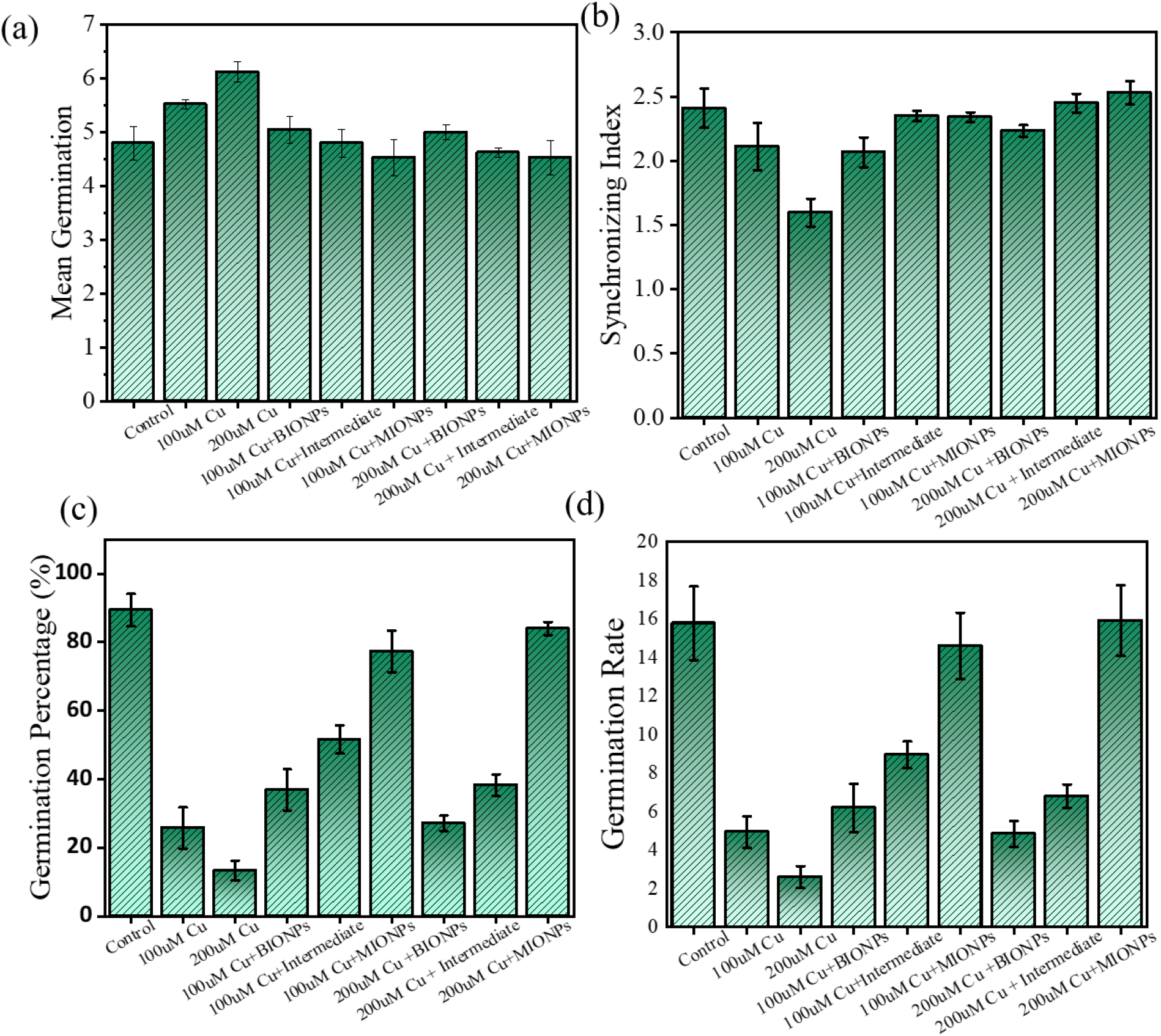
Germination kinetics of Solanum lycopersicum seeds under copper stress and nanoparticle treatment, monitored over 7 days across nine treatment groups (control; 100 µM and 200 µM Cu^2+^; and each Cu^2+^ concentration combined with BIONPs, Intermediate, or MIONPs). (a) Mean germination time (MGT). (b) Synchronization index (SI). (c) Germination percentage (GP). (d) Germination rate (GR). Data are expressed as mean ± SD (n = 3 biological replicates, 75 seeds per treatment).

For insight into the underlying cellular mechanisms of growth inhibition or treatment-mediated recovery, subcellular analyses were subsequently performed. ROS accumulation was examined first because Cu^2+^ induced redox imbalance and excessive ROS generation are early events in cellular stress^47,48^. Excess Cu^2+^ disrupts cellular redox homeostasis and promotes excessive ROS generation, making oxidative stress an important early event in Cu^2+^ -induced phytotoxicity. Therefore, intracellular ROS accumulation was evaluated using H_2_DCFDA fluorescence imaging **(Figure 6)** to determine the extent of oxidative stress induced by Cu^2+^ and the ability of the nanoparticle formulations to restore cellular redox balance. Cu^2+^ -treated seedlings exhibited a pronounced increase in H_2_DCFDA fluorescence **(Figure 5b, c)** compared with the untreated control **(Figure 5a)**, confirming substantial intracellular ROS accumulation. BIONP treatment resulted in a partial reduction in fluorescence **(Figure 5d, g)**, while the intermediate formulation produced a further decrease **(Figure 5e, h)**. MIONP treatment showed the reduction in ROS-associated fluorescence, indicating efficient attenuation of Cu^2+^-induced oxidative stress **(Figure 5f, i)**. The reduction in ROS observed with BIONPs can be attributed to the peroxidase- and catalase-like nanozyme activities of the Fe3O4 core, which facilitate ROS decomposition. The additional reduction observed with the intermediate formulation is associated with the antioxidant and redox-buffering contribution of GSH. In MIONPs, these ROS-regulating effects are complemented by MPA-derived thiol groups that bind excess Cu^2+ 2+^, thereby reducing the availability of Cu^2+^for continued redox-mediated ROS generation. Thus, MIONPs act through complementary mechanisms involving ROS decomposition, redox buffering, and reduction of the Cu^2+^burden, providing greater control of oxidative stress than the individual formulations. This integrated activity is consistent with the stronger physiological recovery observed in **Figures 4 and 5**.

**Figure 6.**
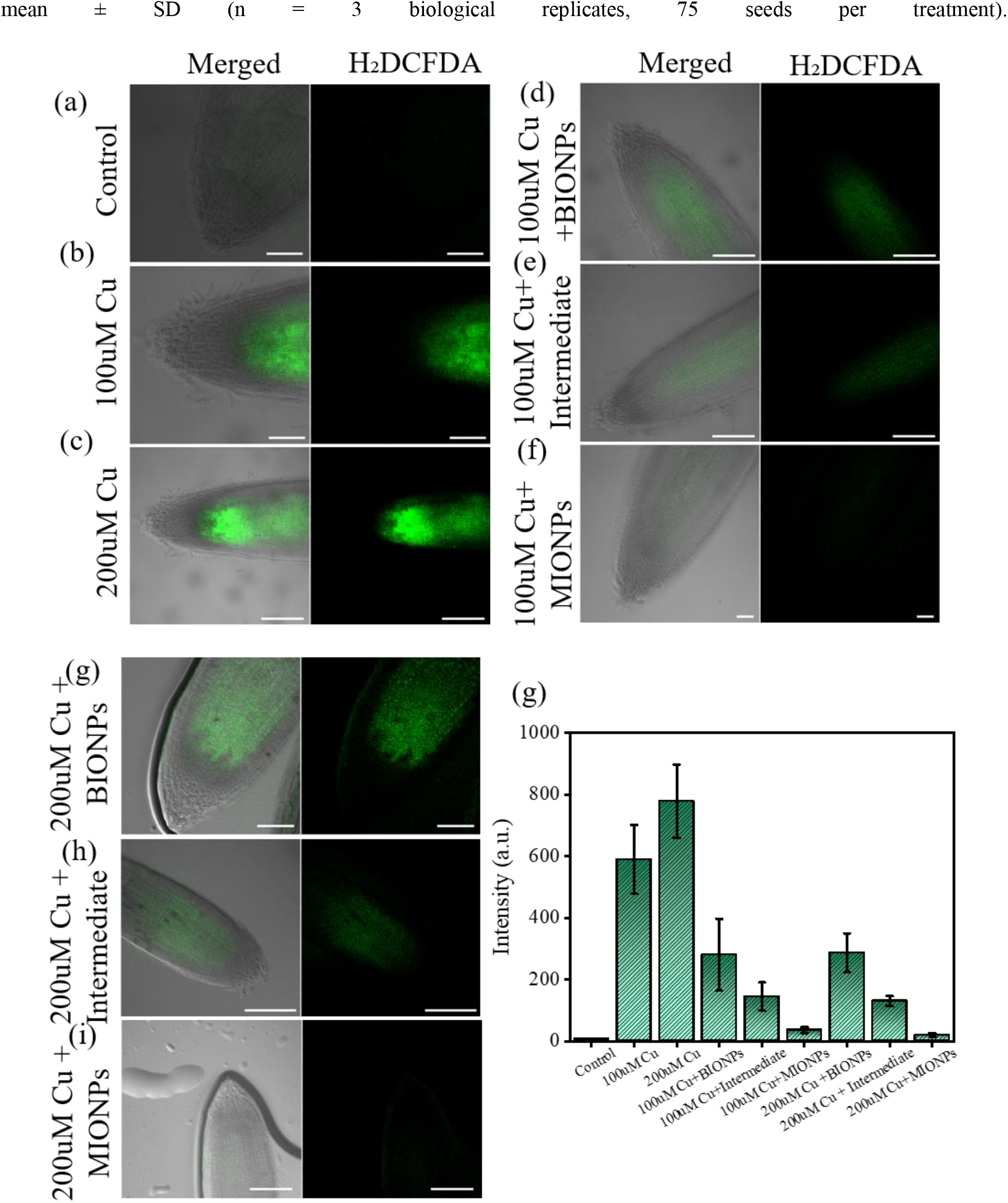
ROS analysis in Solanum lycopersicum seedlings under copper stress and nanoparticle treatment. (I) Representative fluorescence images showing intracellular ROS levels: (a) control, (b) 100 µM Cu^2+^, (c) 200 µM Cu^2+^,(d) 100 µM Cu^2+^ + BIONPs, (e) 100 µM Cu^2+^ + Intermediate, (f) 100 µM Cu^2+^ + MIONPs, (g) 200 µM Cu^2+^ + BIONPs,(h) 200 µM Cu^2+^ + Intermediate, (i) 200 µM Cu^2+^ + MIONPs. (II) Quantitative analysis of fluorescence intensity epresenting relative intracellular ROS levels under different treatments.

The marked reduction in ROS following MIONP treatment indicates attenuation of the Cu^2+^ -induced oxidative stress and restoration of cellular redox homeostasis. Because excessive ROS can impair mitochondrial membranes and disrupt mitochondrial function^49^, the subsequent experiment examined mitochondrial to determine whether the reduction in oxidative stress was accompanied by mitochondrial recovery. Mitochondria are central to cellular energy production and are particularly vulnerable to Cu^2+^-induced oxidative damage. Excess ROS can impair mitochondrial structure and function, making mitochondria an important downstream target of Cu^2+^-induced oxidative stress. Therefore, mitochondrial distribution and integrity were examined using MitoTracker Green fluorescence **(Figure 7)** to determine the extent of mitochondrial alterations under Cu^2+^ stress and to assess whether nanoparticle treatment could preserve mitochondrial organization. Cu^2+^-treated seedlings exhibited a pronounced reduction in fluorescence compared with the untreated control, indicating alterations in mitochondrial organization under Cu^2+^ stress **(Figure 7a, b and c)**. BIONP treatment produced partial restoration of the fluorescence signal **(Figure 7d, g)**, while the intermediate formulation resulted in greater recovery **(Figure 7e, h)**. MIONP treatment showed the strongest restoration of fluorescence, indicating improved preservation of mitochondrial organization under Cu^2+^ stress **(Figure 7f, i)**.

**Figure 7.**
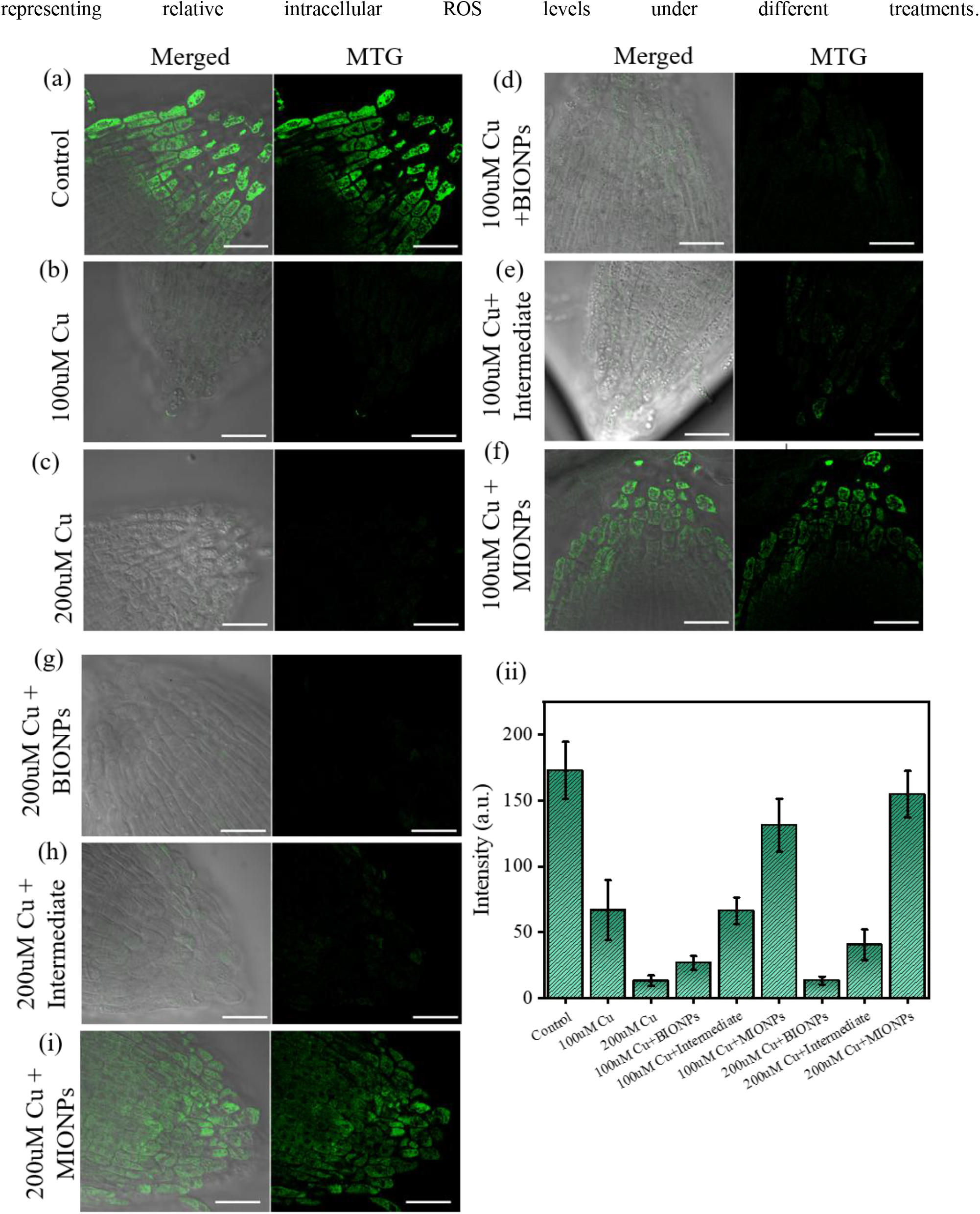
Mitochondrial activity analysis in Solanum lycopersicum seedlings under copper stress and nanoparticletreatment. (I) Representative fluorescence images showing mitochondrial activity: (a) control, (b) 100 µM Cu^2+^,(c) 200 µM Cu^2+^, (d) 100 µM Cu^2+^ + BIONPs, (e) 100 µM Cu^2+^ + Intermediate, (f) 100 µM Cu^2+^ + MIONPs, (g) 200 µM Cu^2+^ + BIONPs, (h) 200 µM Cu^2+^ + Intermediate, (i) 200 µM Cu^2+^ + MIONPs. (II) Quantitative analysis of fluorescence intensity representing relative mitochondrial activity under different treatments.

The mitochondrial recovery observed with MIONPs was associated with attenuation of the Cu^2+^-induced oxidative stress observed in **Figure 6**. MPA-derived thiol groups chelate excess Cu^2+^, reducing its intracellular availability, while the Fe3O4 core contributes nanozyme-mediated ROS decomposition and GSH provides redox buffering. Together, these functions decrease ROS accumulation and restore the cellular redox environment, thereby promoting recovery of mitochondrial organization under Cu^2+^ stress. The stronger mitochondrial recovery with MIONPs compared with BIONPs and the intermediate formulation further supports the contribution of their integrated Cu^2+^-binding and antioxidant functionalities. Because persistent mitochondrial alterations and oxidative stress can extend cellular damage to the nucleus, nuclear chromatin organization was subsequently examined to determine whether MIONP-mediated mitochondrial protection was accompanied by recovery at the nuclear level.

The nucleus represents an important downstream target of Cu^2+^ -induced cellular stress, as persistent oxidative stress can alter nuclear organization and chromatin structure^50^. Excess Cu^2+^ promotes ROS accumulation and mitochondrial dysfunction^51^, which can intensify cellular stress and contribute to chromatin condensation. Therefore, nuclear organization was examined as a downstream indicator of Cu^2+^ -induced cellular damage and to determine whether the protective effect of MIONPs extends to the nuclear level. DAPI fluorescence imaging **(Figure 8)** was used to evaluate nuclear morphology and chromatin organization in Cu^2+^ -stressed seedlings following treatment with BIONPs, the intermediate chitosan/GSH formulation, and MIONPs. Cu^2+^ exposure resulted in pronounced chromatin condensation compared with the untreated control, indicating substantial alteration of nuclear organization under Cu^2+^ stress **(Figure 8a, b and c)**. BIONPs treatment partially reduced chromatin condensation, while the intermediate formulation produced greater recovery **(Figure 8d, g)**. MIONP treatment showed the preservation of nuclear organization, with chromatin distribution approaching that of the control **(Figure 8f, i)**. Quantitative analysis further confirmed a significant reduction in chromatin condensation following MIONP treatment.

**Figure 8.**
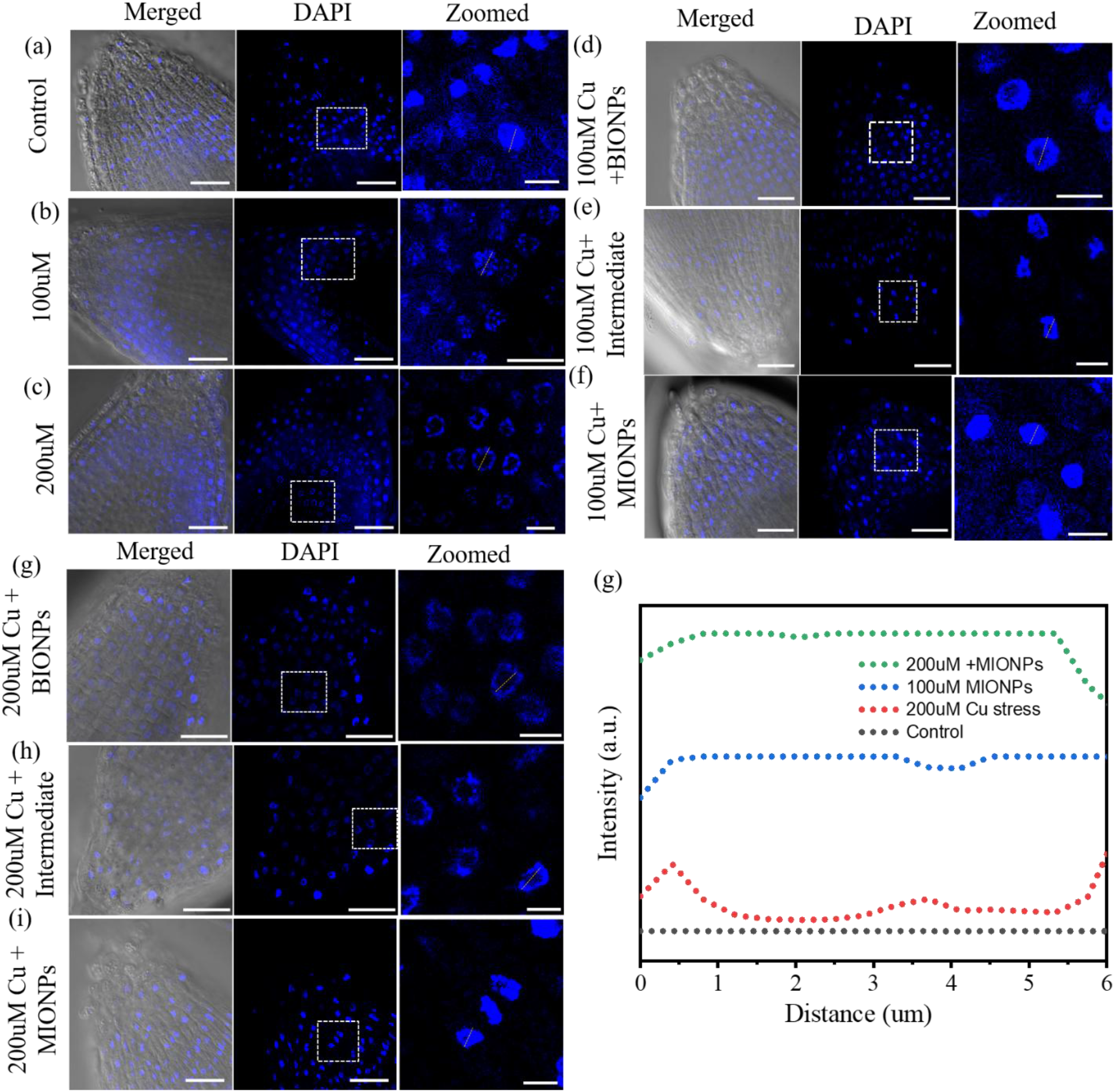
Nuclear morphology analysis using DAPI staining and confocal microscopy in Solanum lycopersicum seedlings. (I) Representative images: (a) control, (b) 100 µM Cu^2+^, (c) 200 µM Cu^2+^, (d) 100 µM Cu^2+^ + BIONPs, (e) 100 µM Cu^2+^ + Intermediate, (f) 100 µM Cu^2+^ + MIONPs, (g) 200 µM Cu^2+^ + BIONPs, (h) 200 µM Cu^2+^ + Intermediate, (i) 200 µM Cu^2+^ + MIONPs. Insets show magnified nuclear structure. (II) Comparative analysis of nuclear fluorescence distribution indicating chromatin organization state.

The recovery of nuclear organization following MIONP treatment can be explained by the reduction of the upstream Cu^2+^ -induced oxidative and mitochondrial stress. MPA-derived thiol groups bind excess Cu^2+ 2+^, reducing its availability for Cu^2+^ -mediated redox reactions and thereby limiting continued ROS generation. At the same time, the Fe3O4 core contributes ROS-regulating nanozyme activity, while GSH supports cellular redox buffering. The resulting decrease in ROS reduces oxidative stress on nuclear components and limits chromatin condensation. In parallel, the reduction in oxidative stress helps preserve mitochondrial membrane potential, thereby limiting further propagation of cellular stress toward the nucleus. Consequently, MIONP treatment promotes recovery of nuclear organization through coordinated reduction of the Cu^2+^burden, attenuation of oxidative stress, and preservation of mitochondrial function.

Overall, the cellular results demonstrate a sequential protective response in which Cu^2+^sequestration reduces the metal-associated oxidative burden, ROS attenuation protects mitochondrial integrity, and preservation of mitochondrial function limits downstream nuclear damage. The resulting recovery of nuclear organization is consistent with the improved germination and root/shoot development observed following MIONP treatment. Thus, MIONPs provide protection against Cu^2+^ stress through coordinated regulation of the Cu^2+^burden, oxidative stress, mitochondrial dysfunction, and nuclear damage, rather than acting solely at a single cellular level. Together, **Figures 4-8** establish a coherent, multi-level mechanistic narrative: Cu^2+^toxicity originates from Fenton-driven ROS generation, which propagates through mitochondrial depolarization to nuclear chromatin damage, ultimately result in impaired germination and growth. Each nanoparticle formulation intervened at a different stage of this cascade, but only MIONPs addressed the problem at its source (Cu^2+^ sequestration) while simultaneously mitigating its downstream consequences (ROS accumulation, mitochondrial dysfunction, and chromatin damage). These findings indicate that the combined Cu^2+^-binding and antioxidant functions of MIONPs provided greater protection than BIONPs and the intermediate nanocomposite across the physiological and cellular parameters examined, highlighting their potential for mitigating Cu^2+^-induced phytotoxicity.

**Scheme 1.**
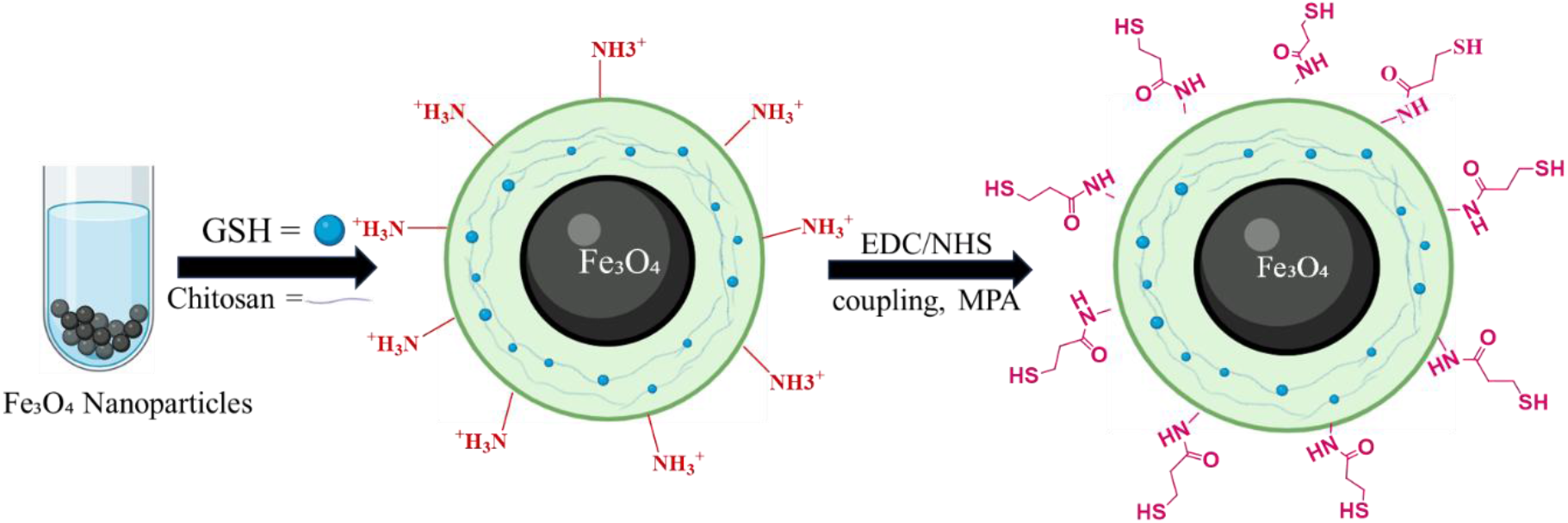
Schematic illustration of the stepwise synthesis and surface functionalization of BIONPs into the chitosan/GSH-coated intermediate and finally MIONPs, functionalized with glutathione (GSH) and 3-mercaptopropionic acid (MPA).

## Conclusions

MIONPs demonstrated strong protective effects against Cu^2+^ stress in *Solanum lycopersicum* seedlings exposed to 100 and 200 µM Cu^2+^. Treatment with MIONPs improved seed germination and promoted recovery of root and shoot growth under Cu^2+^ exposure. At the cellular level, MIONPs reduced intracellular ROS accumulation, restored mitochondrial membrane potential, and preserved nuclear and chromatin organization. Among the tested formulations, MIONPs provided the greatest protection compared with bare BIONPs and intermediate, shows the advantage of their multifunctional design. The combined Fe3O4, GSH, and MPA-derived thiol groups contributed to Cu^2+^ sequestration, redox regulation, and cellular protection. These results indicate that MIONPs can simultaneously reduce Cu^2+^ burden and limit downstream oxidative and organelle damage, thereby supporting plant growth under metal stress. Overall, MIONPs represent a promising nanotechnology-based approach for mitigating Cu^2+^ toxicity, with potential application in protecting crops and managing Cu^2+^-contaminated agricultural soils. Further soil and field studies are required to evaluate their long-term environmental safety and practical applicability. Moreover, the Fe3O4 core provides catalytic oxidase-like activity, while GSH contributes antioxidant protection and chitosan enhances biocompatibility, suggesting that MIONPs may have broader potential for mitigating other stress conditions beyond Cu^2+^ toxicity.

## Supporting information

Supplementary Informations

## Notes

### Competing Interest Statement

The authors have declared no competing interest.

