## Supplementary Informations for "Eifficient Mitigation of Copper Induced Cellular Dysfunction Using Chitosan Based Iron Oxide Nanoparticles"

### Materials and Methods

#### Materials

Iron(III) chloride hexahydrate ($FeCl_3{\cdot}6H_2O$), iron(II) chloride tetrahydrate ($FeCl_2{\cdot}4H_2O$), chitosan (medium molecular weight, degree of deacetylation ≥75%), sodium tripolyphosphate (TPP), glutathione (GSH, reduced form), 3-mercaptopropionic acid (MPA), 1-ethyl-3-(3-dimethylaminopropyl)carbodiimide hydrochloride (EDC), N-hydroxysuccinimide (NHS), copper sulfate pentahydrate ($CuSO_4{\cdot}5H_2O$), 2-(N-morpholino)ethanesulfonic acid (MES buffer), ammonia solution (25%), and acetic acid (glacial) were purchased from Sigma-Aldrich. 2',7'-Dichlorodihydrofluorescein diacetate ($H_2$DCFDA), MitoTracker Red, and 4',6-diamidino-2-phenylindole (DAPI) were obtained from Thermo Fisher Scientific. *Solanum lycopersicum* seeds. All chemicals were of analytical grade and used without further purification. Milli-Q water (resistivity ≥18.2 MΩ·cm) was used throughout.

#### Synthesis of $Fe_3O_4$ Nanoparticles

Bare $Fe_3O_4$ nanoparticles were synthesized by the conventional co-precipitation method following the Massart protocol with minor modifications [36]. Briefly, $FeCl_3{\cdot}6H_2O$ (2.0 mmol) and $FeCl_2{\cdot}4H_2O$ (1.0 mmol) were dissolved in 40 mL of Milli-Q water under continuous magnetic stirring at 80°C in a nitrogen atmosphere to prevent oxidation. Aqueous ammonia solution (25%, v/v) was added dropwise until the pH reached 10 -11, whereupon a stable black precipitate formed instantaneously, confirming successful nucleation and growth of $Fe_3O_4$ cores. The reaction was maintained at 80°C for 30 min. The nanoparticles were collected by magnetic decantation, washed repeatedly with Milli-Q water and ethanol, and redispersed in Milli-Q water. Synthesized $Fe_3O_4$ nanoparticles were stored at 4°C until further use.

#### Chitosan Coating and GSH Encapsulation

Chitosan (0.2% w/v) was dissolved in 1% (v/v) acetic acid overnight at room temperature, and pH was adjusted to 5.5 using 1 M NaOH. $Fe_3O_4$ nanoparticles were dispersed in the chitosan solution (1:5 w/v) under probe sonication for 10 min. Glutathione (GSH, 2 mg/mL) was added and stirred for 1 h. Ionic gelation was induced by dropwise addition of sodium tripolyphosphate (TPP, 0.1% w/v), resulting in stable chitosan-coated nanostructures encapsulating GSH.

Nanoparticles were purified by centrifugation (10,000 rpm, 20 min), washed three times with Milli-Q water, and redispersed to yield $Fe_3O_4$/Chitosan/GSH nanoparticles (Intermediate).

**MPA Conjugation via Carbodiimide Chemistry**

Thiol functionalization was achieved by covalent conjugation of MPA using EDC/NHS carbodiimide chemistry. $Fe_3O_4$/Chitosan/GSH nanoparticles (1 mg/mL in MES buffer, 0.1 M, pH 5.5) were activated with EDC (5 mM) and NHS (5 mM) for 30 min. MPA (10 mM) was added and the reaction continued for 4 h at room temperature. Excess reagents were removed by dialysis (MWCO 12 -14 kDa, 24 h). The purified $Fe_3O_4$/Chitosan/GSH/MPA (MIONPs) nanoparticles were lyophilized and stored at 4°C.

**Physicochemical Characterization**

Morphology and particle size were examined by transmission electron microscopy (TEM). Hydrodynamic size and zeta potential were measured by dynamic light scattering (DLS,) at 25°C. Surface functional groups were characterized by FTIR spectroscopy (ATR mode, 400 - 4000 $cm^{-1}$). Surface elemental composition was analyzed by X-ray photoelectron spectroscopy (XPS). Crystalline phase was confirmed by powder X-ray diffraction (XRD). Iron content at each functionalization stage was quantified by atomic absorption spectroscopy. Copper binding efficiency was determined by inductively coupled plasma mass spectrometry.

**ICP-MS $Cu^{2+}$ Binding Efficiency Assay**

A copper stock solution (1000 mg $L^{-1}$) was prepared by dissolving $CuSO_4 \cdot 5H_2O$ (0.197 g) in 50 mL ultrapure water. Working solutions (0.5 mg $L^{-1}$) were prepared daily by dilution. Adsorption experiments were carried out by adding 0.5 mg of each adsorbent to 5 mL of the 0.5 mg $L^{-1}$ Cu(II) working solution under agitation. After incubation, suspensions were centrifuged and supernatants collected. Prior to ICP-MS measurement, 2.0 mL supernatant was diluted to 10 mL with 2% $HNO_3$ (v/v). Equilibrium adsorption capacity ($Q_e$, µg $g^{-1}$) and percentage binding efficiency were calculated as:

$Q_e = (C_0 - C_e) \times V / m$ (Equation 1)

$\% \text{ Binding} = [(C_0 - C_e) / C_0] \times 100$ (Equation 2)

where $C_0$ (µg $L^{-1}$) is the initial Cu concentration, $C_e$ (µg $L^{-1}$) is the equilibrium concentration, V (L) is the solution volume, and m (g) is the adsorbent mass.

**Seed Germination and Seedling Growth Assay**

*Solanum lycopersicum* seeds were surface-sterilized by sequential immersion in 70% ethanol (1 min), 1% sodium hypochlorite (10 min), and rinsed five times with sterile Milli-Q water. Seeds were germinated on Whatman No. 1 filter paper in Petri dishes (90 mm) under controlled conditions (25 ± 2°C, 16 h light/8 h dark, 150 µmol $m^{-2}$ $s^{-1}$). Nine treatment groups were established: (i) control; (ii) 100 µM $CuSO_4$; (iii) 200 µM $CuSO_4$; (iv) 100 µM Cu + BIONPs;

(v) 200 μM Cu + BIONPs (vi) 100 μM Cu + intermediate (vii) 200 μM Cu + intermediate (viii) 100 μM Cu + MIONPs and (ix) 200 μM Cu + MIONPs. Nanoparticle concentration was 100 μg/mL across all nanoparticle treatments. Germination percentage, root length, and shoot length were recorded after 15 days. Each treatment was performed in triplicate with 20 seeds per Petri dish.

**Germination Kinetics Analysis (GP, GR, MGT, SI)**

To further resolve the temporal dynamics of germination under copper stress and nanoparticle-mediated amelioration, healthy and uniform seeds were surface-sterilized with sodium hypochlorite solution and rinsed repeatedly with distilled water before being placed in Petri dishes across the same nine treatment groups described above (control; 100 μM and 200 μM $CuSO_4$; and each Cu concentration combined with BIONPs, intermediate, or MIONPs. Each treatment comprised 75 seeds divided into three biological replicates. Petri dishes were incubated under controlled laboratory conditions, and the number of newly germinated seeds was recorded daily for each replicate over a 7-day period. The resulting daily germination counts were used to calculate the germination percentage (GP), germination rate (GR), mean germination time (MGT), and synchronization index (SI), as follows:

$GP = (n / N) \times 100$ (Equation 3)

where n is the total number of germinated seeds and N is the total number of seeds sown. GP was used to determine the percentage of successfully germinated seeds under each treatment condition.

$GR = \Sigma (n_i / t_i$ (Equation 4)

where $n_i$ is the number of seeds germinated on day i and $t_i$ is the germination time in days. GR was used to evaluate the speed of seed germination, with higher GR values indicating faster germination.

$MGT = \Sigma(n_i t_i) / \Sigma n_i$ (Equation 5)

where $n_i$ and $t_i$ are as defined above. MGT was used to determine the average time required for seed germination, with lower MGT values indicating faster germination.

$SI = - \Sigma f_i \log_2(f_i)$ (Equation 6)

where $f_i$ is the relative germination frequency on day i. SI was used to assess the uniformity and synchronization of seed germination among treatments. All germination kinetics experiments were performed in triplicate, and data are expressed as mean ± standard deviation (SD); the calculated parameters were plotted using Origin software with error bars representing SD.

**Confocal Microscopy**

All confocal imaging was performed using a Nikon Eclipse Ti inverted confocal microscope (Nikon Instruments), and image acquisition was conducted using Nikon NIS-Elements software. A 60× oil immersion objective (NA 1.40) was used throughout. Four laser lines with corresponding filter sets were employed: 405 nm (DAPI), 488 nm (FITC), 591 nm (TRITC), and 639 nm (CY5). All acquisition parameters detector gain, laser power, pinhole diameter were held strictly constant across all treatment groups. Using NIS-Elements software and displayed with identical brightness and contrast settings across all groups.

**Intracellular ROS Detection Using $H_2$DCFDA**

Intracellular ROS were detected using $H_2$DCFDA (Sigma-Aldrich, D6883), which is deacetylated by intracellular esterases to $H_2$DCFDA and subsequently oxidized by ROS to the highly fluorescent $H_2$DCFDA product [37,38]. A stock solution (10 mM in DMSO) was freshly prepared before use. Root tips (~1 cm) from 15-day-old seedlings were incubated in $H_2$DCFDA working solution (25 µM in PBS, pH 7.4) for 30 min at 25°C in the dark. Root tips were washed three times with PBS (5 min each), hand-sectioned, mounted on poly-L-lysine coated slides, and imaged within 30 min. Controls included: unstained (autofluorescence), positive (200 µM $H_2O_2$), and probe-only controls $H_2$DCFDA fluorescence was excited at 488 nm (FITC channel).

**Mitochondrial Activity Assessment Using MitoTracker Red**

Mitochondrial membrane potential ($\Delta\Psi m$) was assessed using MitoTracker, which accumulates in active mitochondria proportionally to $\Delta\Psi m$. Root tips were incubated in 200 nM MitoTracker Red in half-strength MS medium for 45 min at 25°C in the dark, washed three times (5 min each), sectioned, and mounted immediately. MitoTracker Red fluorescence was excited at 591 nm (TRITC channel).

**Nuclear Morphology Analysis Using DAPI**

Nuclear organization was examined by DAPI staining. Root tip sections were fixed in 4% paraformaldehyde in PBS (pH 7.4) for 45 min, washed three times with PBS, and permeabilized with 0.5% Triton X-100 for 15 min. Sections were stained with DAPI (1 µg/mL in PBS) for 20 min in the dark, washed. DAPI fluorescence was excited at 405 nm. Nuclear fluorescence distribution and chromatin condensation patterns were assessed qualitatively and quantitatively using the NIS-Elements line profile tool.

**Quantitative Image Analysis**

Fluorescence intensity was quantified from maximum intensity projections using NIS-Elements. $H_2$DCFDA (25 µM, 30 min), MitoTracker Red (200 nM, 45 min), and DAPI (1 µg/mL, 20 min) stained sections were imaged separately for each treatment group under identical settings. All values were normalized to the control group and expressed as relative fluorescence intensity ± SD.

**Statistical Analysis**

All experiments were performed with three independent biological replicates ($n = 3$) and data expressed as mean ± SD. Statistical significance was determined by paired t-test. Differences were considered statistically significant at $p < 0.05$.

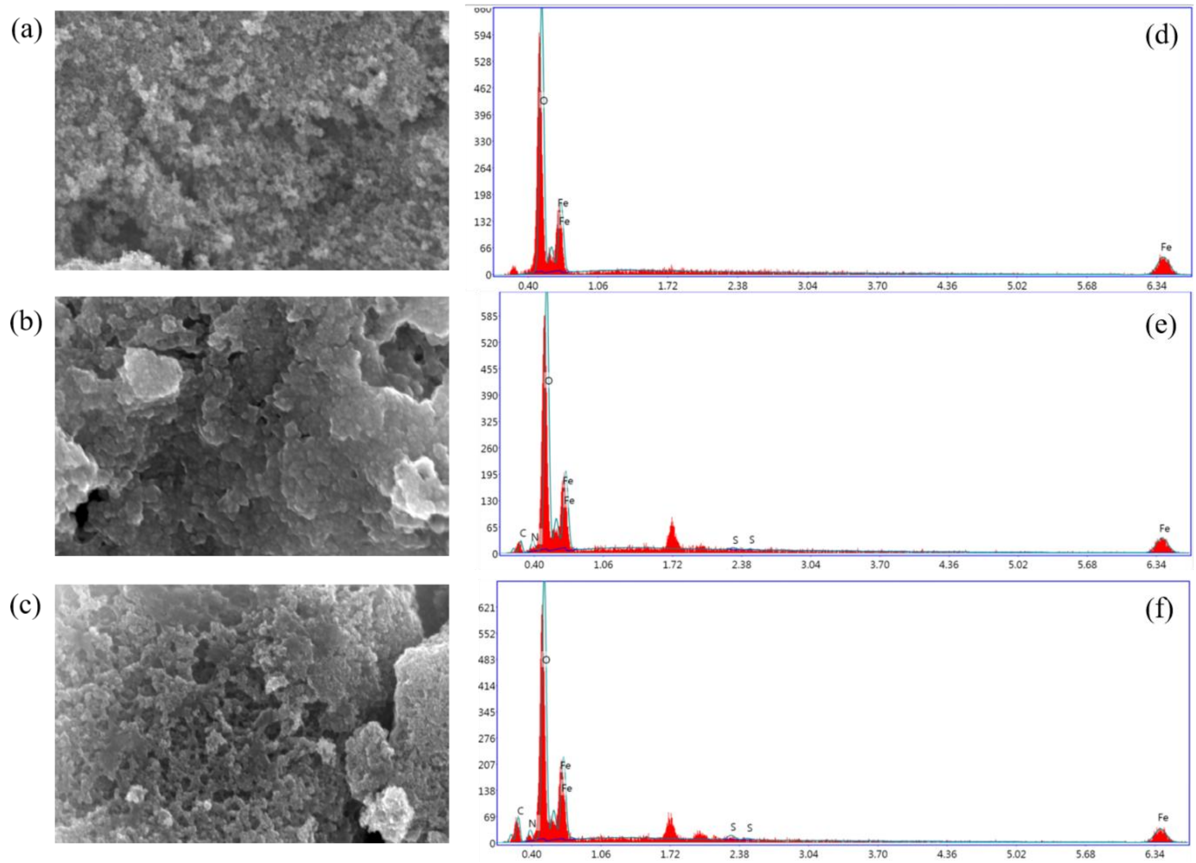


**Figure S1.** SEM images of (a) BIONPs, (b) intermediate, and (c) MIONPs, with (d-e) their corresponding EDX spectra, respectively.

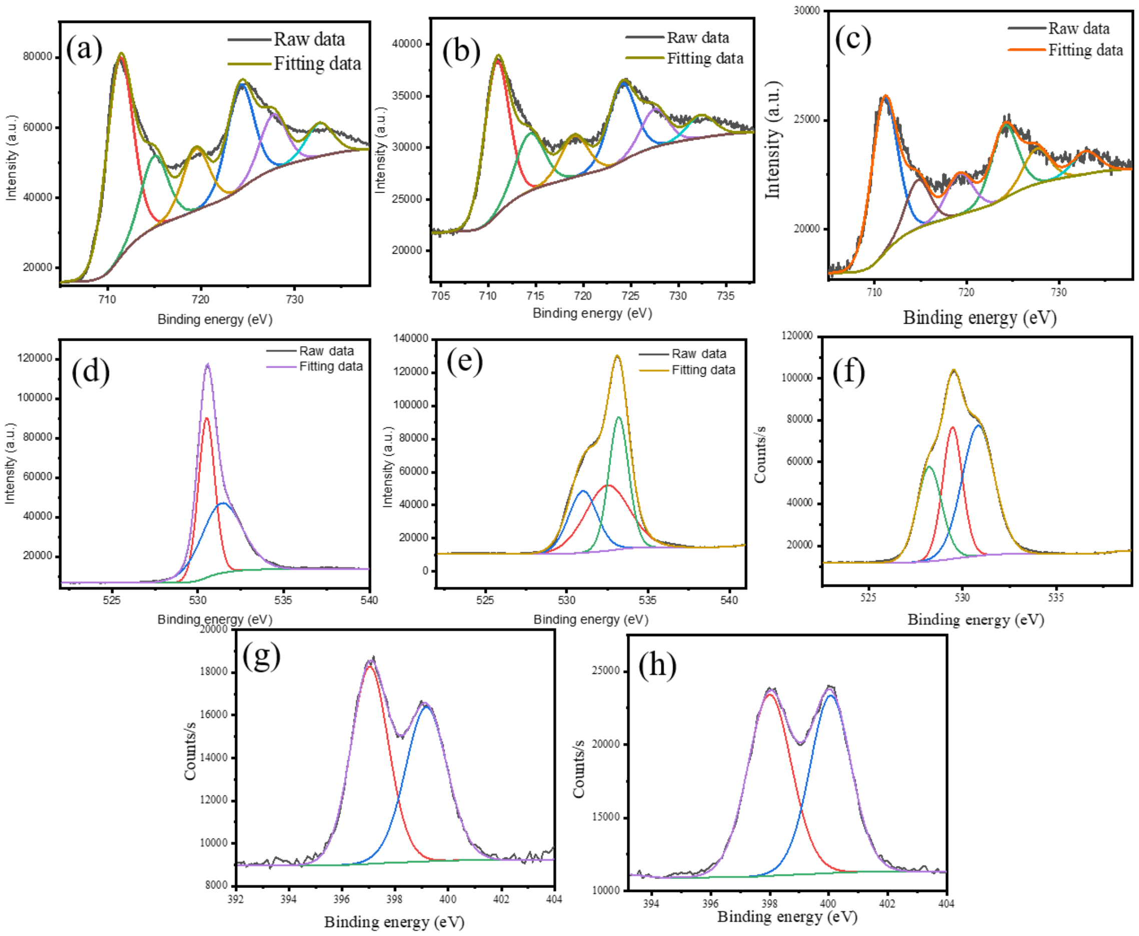


**Figure S2.** XPS deconvolution data of (a) Fe2p of BIONPs, (b-c) Fe2p of the intermediate and MIONPs, respectively; (d-f) O1s of BIONPs, the intermediate, and MIONPs, respectively; (g-h) N1s of the intermediate and MIONPs, respectively.

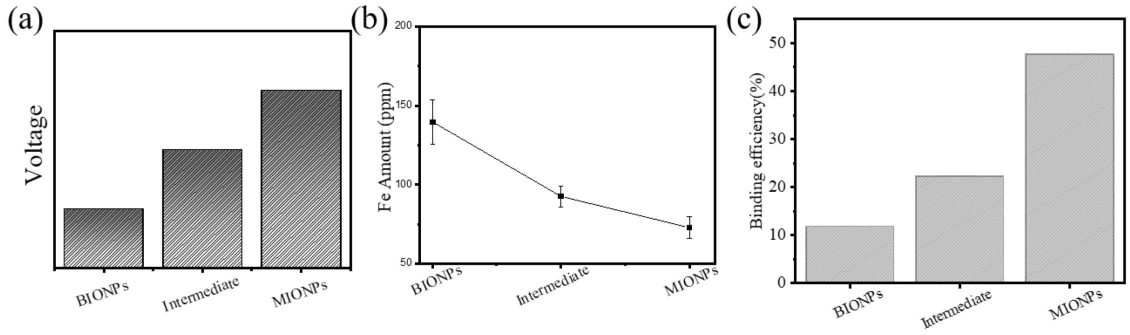


**Figure S3. (**a) Zeta potential of BIONPs, the intermediate, and MIONPs. (b) AAS quantification of Fe content in each material. (c) ICP-MS analysis showing Cu content for BIONPs, the intermediate, and MIONPs.

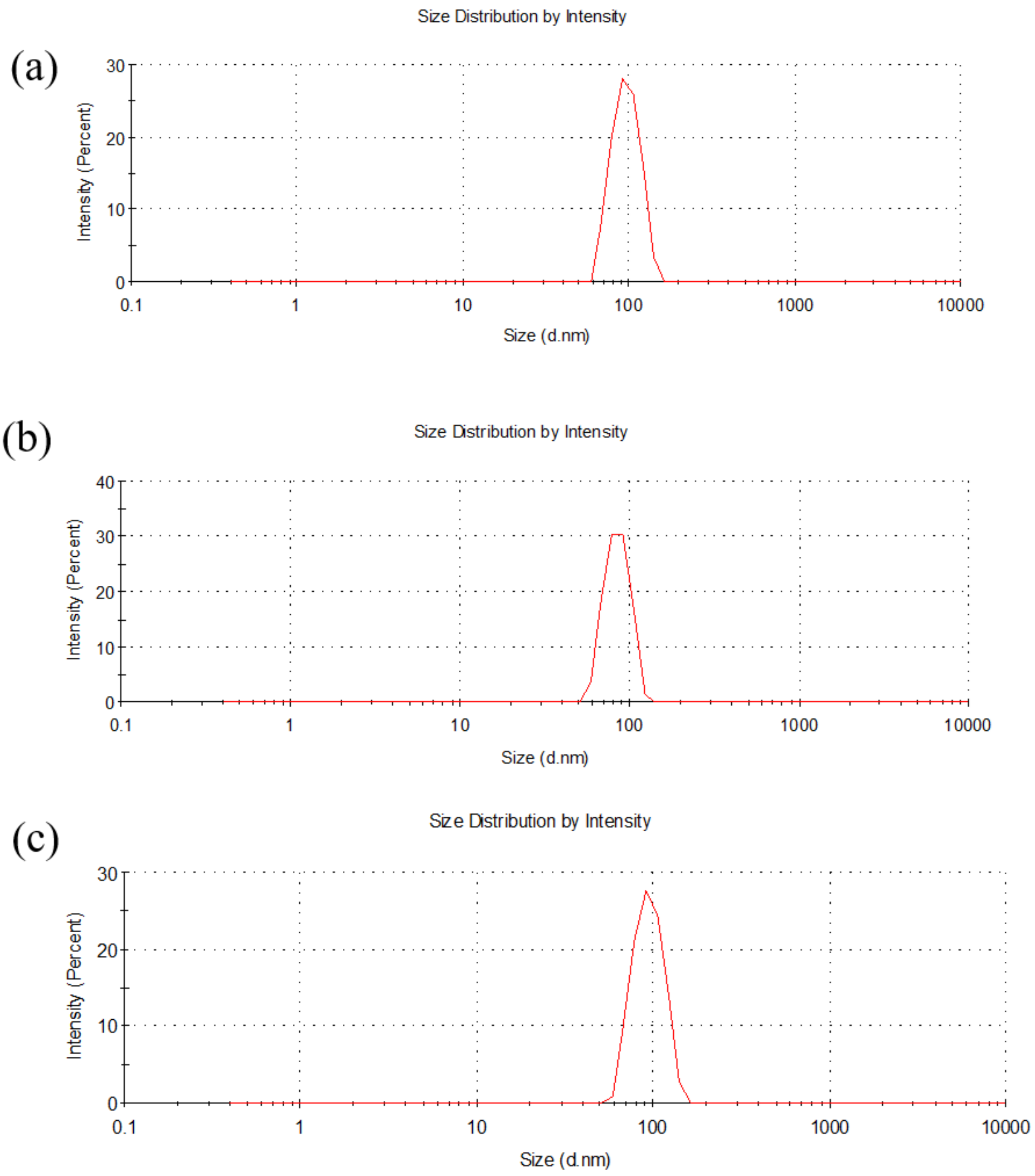


**Figure S4.** Hydrodynamic size distribution profiles obtained by dynamic light scattering (DLS) for (a) BIONPs, (b) the intermediate, and (c) MIONPs, alongside the free chitosan, GSH, and MPA reagents, showing the change in hydrodynamic diameter at each stage of surface functionalization.
